# Methylation-wide association analysis reveals AIM2, DGUOK, GNAI3, and ST14 genes as potential contributors to the Alzheimer’s disease pathogenesis

**DOI:** 10.1101/322503

**Authors:** Alireza Nazarian, Anatoliy I. Yashin, Alexander M. Kulminski

## Abstract

**Introduction:** Alzheimer’s disease (AD) is a progressive complex neurodegenerative disorder with devastating impact on cognitive abilities. It is among the top 10 leading causes of death in the United States with no curative medications. Exploring genetic and non-genetic contributors to AD development is, therefore, of great importance.

**Methods:** We investigated the AD-associated epigenetic changes by combing results from publicly available genome-wide association analyses and a large-scale methylation quantitative trait loci study.

**Results:** Probes mapped to 133 genes were associated with AD with < 2.50E-06. Of these, four genes (i.e., GNAI3, AIM2, DGUOK and ST14) provided stronger evidence of possible role in AD pathogenesis as they were also significantly associated with AD in previous expression quantitative trait loci analyses and/or mouse model studies.

**Discussion:** Although the identified associations do not prove any definitive causal relationships with AD, they provide a list of prioritized genes for follow-up functional studies.

## Introduction

Alzheimer’s disease (AD) is the major cause of dementia worldwide that is projected to affect more than 13 million people in the United States by 2050, with huge health and economic burdens (Alzheimer’s Association 2016; Ridge *et al.* 2016). A rare familial form of AD is inherited as a Mendelian disorder due to autosomal dominant mutations in APP, PSEN1, or PSEN2 genes. Most AD cases, however, are complex diseases arising from the interaction of many genetic and non-genetic factors (Raghavan and Tosto 2017). Thousands of genetic variants in several chromosomal regions and genes have thus far been associated with the common forms of AD (Leslie *et al.* 2014; Ridge *et al.* 2016; Raghavan and Tosto 2017), although the vast majority of cases cannot be etiologically attributed to these variants (Sanchez-Mut and Gräff 2015; Ridge *et al.* 2016; Raghavan and Tosto 2017). Also, none of the non-genetic risk factors, such as age, blood cholesterol and glucose levels, obesity, inactivity, hypertension, head trauma, education level, smoking, alcohol consumption, and depression are proven to have strong causal relationships with AD (Daviglus *et al.* 2010; Power *et al.* 2011; Sanchez-Mut and Gräff 2015).

The epigenetics mechanisms that mainly act in the interaction with non-genetic factors to impact gene expressions were hypothesized to contribute to AD development (Lahiri *et al.* 2008), particularly because of the heterogeneous clinical manifestations observed in subjects with similar or identical genetic backgrounds (Yokoyama *et al.* 2017). The potential impact of epigenetic changes on AD pathogenesis has been widely investigated in cell lines, mouse models, and post-mortem brain samples. These studies provided evidence of association of epigenetic changes with AD, albeit the detected differences were sometimes inconclusive and non-replicated (Sanchez-Mut and Gräff 2015; Yokoyama *et al.* 2017). Besides several candidate-gene studies that reported changes in methylation profile of some well-known AD genes such as APP, MAPT (Iwata *et al.* 2014), and APOE (Wang *et al.* 2008); genome-wide methylation analyses also identified a number of epigenetically deregulated AD-associated genes. While in most cases such associations were uniquely found in a single study (Sanchez-Mut and Gräff 2015; Yokoyama *et al.* 2017), the AD-associated epigenetic modifications in several genes including ANK1, CDH23, RHBDF2, RPL13-C10orf54 (De Jager *et al.* 2014; Lord and Cruchaga 2014; Lunnon *et al.* 2014), and SORB3 (Siegmund *et al.* 2007; Sanchez-Mut *et al.* 2013) were replicated in independent studies. Also, Yu et al. (2015) reported that ABCA7, BIN1, HLA-DRB5, SLC2A4, and SORL1 genes, that had already been associated with AD in previous genome-wide association studies (GWAS), were also associated with AD epigenetically (Yu *et al.* 2015).

To better understand the mechanisms underlying AD pathogenesis, we investigated the AD-associated epigenetic changes using summary data-based Mendelian randomization (SMR) method (Zhu *et al.* 2016). This method perform methylation-wide association analysis (MWA) of any trait of interest by integrating summary results from a genome-wide association analysis (GWA) of the trait with data from a methylation quantitative trait loci (mQTLs) study. We used summary results from our recently conducted GWA meta-analyses of AD [Nazarian et al. (2018) - manuscript in review] along with data from a large-scale mQTLs study of methylation profile of blood cells (McRae *et al.* 2017) to obtain a set of epigenetically AD-associated genes. We further validated the significant findings by comparing our MWA results with those from our previous transcriptome-wide association analysis (TWA) of AD [Nazarian et al. (2018) - manuscript in review] that implemented the SMR method using the same GWA summary results along with data from an expression quantitative trait loci (eQTLs) study of blood cells by (Lloyd-Jones *et al.* 2017).

## Methods

### Study Participants

This study makes use of the results from our previous GWA meta-analyses [Nazarian et al. (2018) - manuscript in review]. Briefly, these meta-analyses combined the results from GWA analyses of AD susceptibility in four independent cohorts, including: 1) Late-Onset Alzheimer’s Disease Family Study from the National Institute on Aging (NIA-LOADFS) (Lee *et al.* 2008), 2) Framingham Heart Study (FHS) (Dawber *et al.* 1951; Feinleib *et al.* 1975), 3) Cardiovascular Health Study (CHS) (Fried *et al.* 1991), and 4) Health and Retirement Study (HRS) (Sonnega *et al.* 2014). The LOADFS and FHS cohorts were family-based studies, whereas the CHS and HRS had population-based design. All four studies underwent thorough institutional review boards (IRBs) reviews and were conducted according to IRB approved guidelines. The subjects with AD were identified either based on the case/control status directly provided by the LOADFS and FHS datasets, or by using International Classification of Disease codes, Ninth revision (ICD-9) for the CHS and HRS datasets. The meta-analyses were performed under five plans in which either: 1)-the entire subjects in each dataset, 2)-only males, 3)-only females, 4)-only subjects with history of hypertension, or 5)-only subjects with no history of hypertension were analyzed. Table 1 contains the number of participants used for these meta-analyses.

**Table 1.** Numbers of cases and controls used in the meta-analyses.

| Analysis Plan | Total | Cases | Controls |
| --- | --- | --- | --- |
| Plan 1: Males and Females | 17480 | 2741 | 14739 |
| Plan 2: Only Males | 7289 | 952 | 6337 |
| Plan 3: Only Females | 10191 | 1789 | 8402 |
| Plan 4: Hypertension-positive Subjects | 10870 | 1262 | 9608 |
| Plan 5: Hypertension-negative Subjects | 4806 | 796 | 4010 |

### GWA Meta-analysis

Detailed information about genetic analyses in the four cohorts of interest and the subsequent meta-analyses can be found in the original publications [Nazarian et al. (2018) - manuscript in review]. Briefly, the GWA analysis in each cohort was performed to detect associations of ^~^2 million genotyped and imputed single-nucleotide polymorphisms (SNPs) with AD by fitting logistic regression (CHS and HRS cohorts) or generalized mixed logistic regression (LOADFS and FHS cohorts) models. The top 2–5 principal components (PCs), and birth cohort and sex (except plans 2 and 3) of subjects were included in the models as fixed effect covariates. In the case of LOADFS and FHS cohorts with family-based design, the family IDs were considered as a random effect covariate in the models. The individual GWA results from four datasets were then combined by a conventional inverse variance meta-analysis.

### MWA analysis

MWA analysis was performed by SMR package (Zhu *et al.* 2016) combining the meta-analysis results of each plan of the study with the publicly available summary results from a mQTLs study (McRae *et al.* 2017). Probes with at least one significant mQTL (i.e., a given SNP with P_mQTL_ < 5E-08) in McRae et al. (2017) study (McRae *et al.* 2017) which was also among the SNPs in conducted meta-analyses were included in our MWA analyses. This resulted in a set of up to 90362 probes with significant cis-mQTLs for the MWA analyses under the five aforementioned plans. The MWA analysis first performs a SMR test to determine the significance of association between any given probe and AD. The resulting P_SMR_ values were ranked and interpreted at an appropriate false discovery rate (FDR) level to account for multiple testing issue following Benjamini and Hochberg (1995) (Benjamini and Hochberg 1995). For each analysis plan, the appropriate FDR threshold was the largest significance level at which the number of possible false-positives among significant probes was less than 1. Probes with significant P_SMR_ at FDR level of 0.001 (analysis plans 2 and 5) or 0.01 (analysis plans 1, 3 and 4) were then selected for the heterogeneity test (i.e., HEIDI test) to determine if a significant association detected between a given probe and AD is likely to arise from the pleiotropic effects of a single locus on both methylation changes and AD status. A significant HEIDI test (P_HEIDI_ < 0.05) rules out pleiotropy (or causality) possibility and, instead, indicates that the detected probe-AD association is a result of linkage between adjacent variants, each affecting AD susceptibility and methylation pattern separately (Zhu *et al.* 2016).

The probes/genes that passed both SMR (i.e., had significant P_SMR_ at considered FDR thresholds) and HEIDI (i.e., P_HEIDI_ ≥ 0.05) tests were considered as novel AD candidate genes if there were no AD-associated SNPs with significant P_GWAS_ at genome-wide significance level (i.e., P_GWAS_ < 5E-08) within their 1 Mb flanking regions in our meta-analyses [Nazarian et al. (2018) - manuscript in review] or in other GWAS studies that have already been uploaded to the GRASP catalog (Leslie *et al.* 2014). Finally, the lists of AD-associated genes from MWA analyses were compared to those resulted from our previous TWA analyses [Nazarian et al. (2018) - manuscript in review] to find out if there is any overlap between epigenetically and transcriptionally AD-associated genes.

## Results

The Supplementary File 1 (Tables S1-S5) contains the results of our MWA analyses in details. We found that 12, 14, 25, 22 and 104 probes passed SMR test at the aforementioned FDR levels (with P_SMR_ between 2.49E-187 and 2.50E-06) and HEIDI test (P_HEIDI_ ≥ 0.05) under analysis plans 1-5, respectively. These probes were mapped to 7, 11, 23, 16 and 76 genes, respectively as in some cases several significant probes were related to a single gene. The top mQTLs corresponding to these significant probes were nominally significant (1.34E-05 ≤ P_GWAS_ ≤ 4.71E-02) in our meta-analyses [Nazarian et al. (2018) - manuscript in review]. None of the probes/genes that were significant in males were significant in females. Also, among the genes that were significant in subjects with hypertension, only ALDH3A1 was significant in hypertension-negative subjects as well.

To determine whether the significantly detected genes can be considered as potentially novel candidates for AD, the associations of SNPs located in their 1 Mb flanking regions with AD were searched. We noticed that there were AD-associated SNPs with P_GWAS_ < 5E-08 within 1 Mb of cg05206559 probe corresponding to NANOS2 gene on chromosome 19q13 in our meta-analyses [Nazarian et al. (2018) - manuscript in review] and the other GWAS studies (Leslie *et al.* 2014). No SNPs with P_GWAS_ < 5E-08 were found in regions around other probes in our meta-analyses. This might be due to the lack of power of these analyses (e.g. due to insufficient sample sizes) (Zhu *et al.* 2016) or the heterogeneity of association signals detected in GWA analyses of the four cohorts (i.e., LOADFS, FHS, CHS, and HRS) that were used for these meta-analyses. However, in addition to NANOS2, AD-associated SNPs with P_GWAS_ < 5E-08 were found within 1 Mb flanking regions of 7 other probes in previous GWAS studies (Leslie *et al.* 2014). These probes (i.e., cg12710176, cg17430214, cg03942922, cg04035728, cg15638306, cg25674938 and cg04698472) were all significant in subjects with no history of hypertension (i.e., analysis plan 5) and were mapped to YOD1, TREM1, NFYA, CHRNA2, NGFR, MUM1 and SIGLEC12 genes.

Investigating the overlaps between MWA and TWA results, we found that among the potential epigenetically AD-associated genes (Supplementary File 1), four genes had significant AD-associated probes in our TWA analyses [Nazarian et al. (2018) - manuscript in review] as well. The significant TWA probes were mapped to GNAI3 which was significant in females (P_SMR_ = 4.00E-06), and AIM2, DGUOK and ST14 which were significant in subjects with no history of hypertension (P_SMR_ = 1.82E-05, 2.18E-05, and 7.78E-07, respectively). Table 2 summarizes MWA results for these four genes.

**Table 2.** Summary statistics for GNAI3, AIM2, DGUOK and ST14 genes from MWA analysis.

| Probe ID | Gene | b<sub>SMR</sub> | SE<sub>SMR</sub> | P<sub>SMR</sub> | P<sub>HEIDI</sub> |
| --- | --- | --- | --- | --- | --- |
| **Plan 3 - Only Females** |  |  |  |  |  |
| cg08644463 | GNAI3 | -0.291 | 0.061 | **1.62E-06** | 2.09E-01 |
| **Plan 5 - Hypertension-negative Subjects** |  |  |  |  |  |
| cg17515347 | AIM2 | -0.418 | 0.087 | **1.43E-06** | 3.42E-01 |
| cg11003133 | AIM2 | -0.462 | 0.097 | **1.86E-06** | 1.73E-01 |
| cg17217296 | AIM2 | -0.210 | 0.043 | **8.79E-07** | 2.44E-01 |
| cg03063511 | DGUOK | -0.247 | 0.041 | **2.74E-09** | 2.44E-01 |
| cg02850715 | ST14 | -0.812 | 0.138 | **4.14E-09** | 7.39E-01 |
| cg21029769 | ST14 | -1.006 | 0.184 | **4.58E-08** | 9.28E-01 |
| cg00496004 | ST14 | -1.322 | 0.277 | **1.84E-06** | 2.77E-01 |

## Discussion

Despite detecting many genetic variants and identifying several non-genetic factor that may play roles in AD susceptibility, the definitive underlying mechanisms in many AD cases is unclear. It has been suggested that epigenetic mechanisms may notably contribute to the heterogeneous nature of AD (Yokoyama *et al.* 2017). To integrate the effects of genetic and non-genetic contributors on AD pathogenesis, Lahiri et al. (2008) suggested the Latent Early life Associated Regulation (LEARn) model (Lahiri *et al.* 2008). According to the this model, exposure to certain environmental factors early in life may result in epigenetic changes with long-lasting effects that alter the expression profile of particular genes years later, which in turn may have phenotypic consequences. The potential role of epigenetic changes in AD pathogenesis has been emphasized in recent years and a growing body of research has emerged to investigate the matter.

In this study, we combined the results from our previous GWA meta-analyses [Nazarian et al. (2018) - manuscript in review] with data from a publicly available mQTLs study of blood cells (McRae *et al.* 2017) to identify potential genes that might be associated with AD epigenetically. Although the pattern of DNA methylation can be tissue-or cell-specific (Smith *et al.* 2014; Sanchez-Mut and Gräff 2015), using mQTLs data from blood is more feasible than brain-specific mQTLs data as determining methylation profile in brain tissue of living humans is not readily available and, in addition, animal models may not demonstrate the same pathologic changes occurring in human (Smith *et al.* 2014). Using data from blood studies may also provide the analyses with more power due to the larger sample sizes (Zhu *et al.* 2016). Several previous studies have shown the utility of blood samples for investigating the AD-associated epigenetic modifications by reporting global or gene-specific methylation changes in AD subjects compared with matched healthy controls (Bollati *et al.* 2011; Chang *et al.* 2014; Di Francesco *et al.* 2015; Nagata *et al.* 2015; Ji *et al.* 2015).

Our results demonstrated epigenetically significant associations of 133 genes with AD under the five analysis plans of interest, which were mostly plan-specific. Of these, the associations of eight genes (i.e., NANOS2, YOD1, TREM1, NFYA, CHRNA2, NGFR, MUM1 and SIGLEC12) with AD were not considered novel as there were AD-associated SNPs with p < 5E-08 within their 1 Mb up-/downstream regions in our [Nazarian et al. (2018) - manuscript in review] or other studies (Leslie *et al.* 2014). The other 125 genes can, therefore, be considered as potentially novel candidate genes for AD. However, worthy of mention is that the identified associations between these genes and AD do not prove any definitive causal relationships. Instead, they provide a list of prioritized genes for further functional studies (Zhu *et al.* 2016). Such follow-up studies are needed to validate their potential roles in AD pathogenesis.

Epigenetic modifications were suggested to contribute to AD pathogenesis by regulating genes expression through modifying chromatin structure and accessibility (Sanchez-Mut and Gräff 2015). For instance, Iwata et al. (2014) reported that the changes in methylation pattern of APP and MAPT genes in the temporal lobe of subjects with sporadic AD would impact their expression, which, in turn, may be the underlying mechanism for AD development (Iwata *et al.* 2014). Therefore, comparing the changes in methylome and transcriptome profiles may provide a useful validating tool for further prioritizing the list of significant genes by weeding less relevant and consistent ones. In a previous study, Hannon et al. (2017) were able to detect overlapping mQTL and eQTL signals with potential functional implications for Crohn’s disease and migraine by comparing their SMR-based MWA and TWA analyses (Hannon *et al.* 2017). Also, useful insights about biological mechanisms of AD that can be obtained from empirical gene/protein expression studies (e.g., mouse model studies) may serve as another layer of validation.

We compared the list of epigenetically AD-associated genes from our MWA with transcriptionally AD-associated ones from previously conducted TWA analyses [Nazarian et al. (2018) - manuscript in review] to provide further support for validating the significant findings. These comparisons resulted in a short list of four potential AD-associated genes that had significant probes in both MWA and TWA analyses (i.e., GNAI3 in females and AIM2, DGUOK and ST14 in hypertension-negative subjects with P_SMR_ between 1.86E-06 and 2.74E-09 in MWA analyses and between 2.18E-05 and 7.78E-07 in TWA analyses). AD-associated SNPs with P_GWAS_ < 5E-08 were not found within 1 Mb these genes, although several SNPs with P_GWAS_ < 5E-06 were detected in regions around AIM2 by both our meta-analyses [Nazarian et al. (2018) - manuscript in review] and the other groups (Li *et al.* 2008; Leslie *et al.* 2014).

The review of literature related to genome-wide or candidate gene studies provided additional insights on the potential roles of these 4 genes in AD. *GNAI3* encodes a G-protein functioning as a signal transducer in various cellular signaling cascades. SNPs mapped to GNAI3 were associated with total and low-density lipoprotein (LDL) cholesterol levels (Teslovich *et al.* 2010), and major depression (Shi *et al.* 2011) at suggestive level of associations (i.e., P_GWAS_ < 5E-06) in previous GWAS studies. The elevated blood cholesterol and depression are among AD risk factor (Daviglus *et al.* 2010). Also, Neuner et al. (2017) reported that the protein encoded by mouse ortholog of GNAI3 was overexpressed (with 1.96 fold-change) in hippocampus of AD intact mice compared to 5XFAD transgenic ones (i.e., AD impaired mice) which harbor mutant APP and PSEN1 genes (Neuner *et al.* 2017).

*AIM2* encodes a protein involved in regulating cell proliferation as a tumor suppressor, innate immune system responses and autoimmune diseases (Man *et al.* 2016). AIM2 may initiate inflammasomes formation in response to stimuli such as viruses, bacteria, and damaged cells. Inflammasomes in turn mediate the, maturation and release of pro-inflammatory cytokines such as IL-1β and IL18 that are likely to be involved in AD development (Liu and Chan 2014; Ahn *et al.* 2017). IL-1β can activate astrocytes and microglia cells, and stimulate the release of amyloid-β precursor protein (APP) and amyloid-β (Aβ) from neurons. It may increase in the blood, cerebrospinal fluid (CSF), and brain of AD patients. IL-18 is over-expressed in astrocytes, microglia, and neurons in Aβ plaques. It can increase the Aβ formation and mediate the tau protein hyper-phosphorylation. Its blood level may increase in earlier stages of AD (Liu and Chan 2014). Oz et al. (2009) reported that methylene blue (MB), an inhibitor of inflammasome proteins such as AIM2, NLRP3, and NLRC4 (Ahn *et al.* 2017), may decelerate the production of Aβ plaques and neurofibrillary tangles. MB-based medications were, therefore, suggested as potential treatments for AD (Oz *et al.* 2009). In addition, Wu et al. (2016) reported that AIM2 knockout mouse models demonstrated behavioral changes and impaired auditory fear memory (Wu *et al.* 2016).

*DGUOK* encodes a protein involved in mitochondrial purine metabolism pathway. Mitochondrial dysfunction is an important finding in the neurons of AD patients (Querfurth and LaFerla 2010). SNPs within this gene were not associated with any traits with P_GWAS_ < 5E-06 in previous studies (Leslie *et al.* 2014). In However, Ansoleaga et al. (2015) reported that DGUOK was downregulated in entorhinal and precuneus cortex of patients with AD of stages V-VI and III-IV, respectively, compared to well-matched healthy controls (Ansoleaga *et al.* 2015)

*ST14* encodes a membrane serine protease with tumor suppressor activity. SNPs mapped to ST14 were not associated with any traits with P_GWAS_ < 5E-06 in previous studies (Leslie *et al.* 2014). However, Wirz et al. (2013) noticed that the ortholog of ST14 was overexpressed (with 5.39 fold-changes) in the frontal cortex of APPswe/PS1dE9 transgenic mice, harboring APP and PSEN1 mutants, in response to the development of Aβ plaques (Wirz *et al.* 2013). Also, Yin et al. (2017) reported that the mouse ortholog of ST14 was upregulated in Aβ plaques-associated microglia cells in 5XFAD transgenic mice models compared to AD intact ones (Yin *et al.* 2017).

In conclusion, our MWA analyses revealed a list of 133 epigenetically AD-associated genes, 125 of which were considered potential novel AD candidates. Although these significant associations do not implicate any casualty, they can be used to prioritize genes for functional studies. Comparing our MWA results with evidence from previous TWA analyses, we found out four genes (i.e., GNAI3, AIM2, DGUOK and ST14) provided stronger evidence of possible role in AD pathogenesis as they had significant AD-associated probes in TWA analyses as well, and in addition, there was empirical evidence of their association with AD. The combination of MWA and TWA results, and additional information from empirical studies yielded more consistent results and helped to further prioritize the list of AD-associated genes for follow-up studies.

## Acknowledgments

This research was supported by Grants from the National Institute on Aging (P01AG043352 and R01AG047310). The funders had no role in study design, data collection and analysis, decision to publish, or manuscript preparation. The content is solely the responsibility of the authors and does not necessarily represent the official views of the National Institutes of Health.

For information about LOADFS, FHS, CHS and HRS datasets that were used by meta-analyses under consideration, please see the Supplementary Acknowledgment File.

## Supplementary information

Supplementary File 1 (Tables S1-S5) and Supplementary Acknowledgment File

